# Rapid neural analysis of linguistic stress and meter in continuous speech

**DOI:** 10.64898/2026.05.04.722740

**Authors:** Camila Zugarramurdi, Eleonora Beier, Katsuaki Kojima, Stephanie Powell, Jonathan Liu, Kristin Davis, Kylie Korsnack, Brett Myers, Miriam Lense, Srishti Nayak, Reyna L. Gordon, Cyrille Magne, Yulia Oganian

**Affiliations:** Instituto de Fundamentos y Métodos, Facultad de Psicología, Universidad de la República; Center for Mind and Brain, University of California, Davis; Perinatal Institute, Cincinnati Children’s Hospital Medical Center, Department of Pediatrics, University of Cincinnati College of Medicine; Literacy Studies PhD Program, Middle Tennessee State University; Dept. of Otolaryngology - Head & Neck Surgery, Vanderbilt University Medical Center; Faculty Hub, University of Richmond; Dept. of Communication Sciences and Disorders, University of Utah; Psychology Department, Middle Tennessee State University; Center for Integrative Neuroscience, University of Tübingen

**Keywords:** speech, lexical stress, prosody, EEG, speech perception, speech comprehension, envelope tracking

## Abstract

Continuous speech evolves around vowels, the centerpieces of individual syllables. Vowels vary in linguistic and acoustic salience: Linguistically, stressed syllables are more salient than unstressed syllables: Stress patterns convey critical lexico-semantic and prosodic information, and their regularity defines the speech meter. Acoustically, English vowel intensity cues lexical stress but also marks salient syllables irrespective of stress status. Recent evidence demonstrates rapid neural analysis of vowel intensity and identity during perception of continuous speech. Here, we probe how these processes integrate lexical stress and metrical regularity. We recorded EEG while participants (n=26) listened to children’s stories with either an irregular, speech-like meter, or a regular poetic meter. Stress and meter modulated cortical encoding of vowels throughout processing: Preparatory activity preceded vowel onsets in an irregular meter only, and early sensory responses were enhanced for unstressed vowels, suggesting additional resource allocation during processing of uncertain and less discriminable speech sounds. In contrast, later processing (300-500ms) was stronger for stressed syllables and in irregular meters, suggesting a combined effect of uncertainty and informational content. Finally, responses were stronger for small intensity rises within metrically predicted stressed vowels than in all other conditions. In the time-frequency domain, the spectral profile of neural phase-locking corresponded to spectral signatures of individual evoked responses, syllable and stress rates in the stimuli. Overall, our findings reveal rapid neural integration of stress and metrical expectations in neural processing of continuous speech. These dynamics may underlie the perceptual benefits of metrically regular speech, such as poetry and song lyrics.

## Introduction

Verbal communication hinges on the ability to extract meaning from a continuous and rapidly varying speech signal. However, the information content of speech is not uniform in time. Rather, speech is organized in syllabic units of varying informational content and acoustic salience. Lexically stressed syllables are more informative than unstressed syllables, especially with regard to vowel identity (Altman & Carter, 1989; McAllister, 1991). Furthermore, the alternation of stressed and unstressed syllables defines the speech meter. In regular meters, this alternation is predictable. Such regularities are not only aesthetically pleasing and form the basis for poetry, but also provide large-scale contextual predictions of the stress status, and thus informational content, of individual vowels (Beier & Ferreira, 2018).

Multiple lines of evidence show that lexical stress is an integral part of the speech signal. For instance, lexical stress supports recognition of individual speech sounds (Slowiaczek, 1990; van Donselaar et al., 2005). To accommodate such effects, it has been suggested that stress and phonemes are rapidly recognized in parallel and independently of each other during perception (Reinisch et al., 2010; Schild et al., 2014). Metrical regularity was also found beneficial for further integration of speech content, including syllable segmentation (Nazzi & Ramus, 2003) and lexico-semantic access (Rothermich et al., 2012; Zora et al., 2023). Moreover, deviations from metrical patterns are remembered better, providing a prosodic means of emphasis (Kimball, 2018). A large body of electrophysiology studies using M/EEG have found phase-locking between neural signals in auditory cortices and the speech signal in the range of 1 -10 Hz (Boucher et al., 2019; Kolozsvári et al., 2021; Lizarazu et al., 2019; Peelle et al., 2013). This frequency range covers the typical rate at which syllables and stress occur in speech. It was thus hypothesized that this effect reflects neural tracking of syllables, with stronger neural responses to stressed syllables (Gross et al., 2013). However, these analyses typically aggregate across the entire speech signal. It is thus not clear whether phase-locking at syllable and stress rates is driven by stressed or unstressed syllables. More generally, it is not known how this effect may be affected by metrical predictability.

Acoustically, English lexical stress is indexed by the combination of multiple cues, realized predominantly on the vowel nucleus, most importantly vowel intensity (Kandylaki et al., 2022). That is, English stressed vowels are louder and longer than unstressed vowels (Fry, 1955, 1958; Lehiste, 1971; Van Santen, 1992). Critically, vowel intensity is not only overall higher for stressed syllables but also serves as a bottom-up acoustic cue to acoustic salience within each stress category. A reliable acoustic correlate of vowel intensity are rising edges in the speech envelope (“peakRate” events). Recent electrophysiology work revealed that dedicated neural populations in the human speech cortex encode peakRate events and their magnitude, independently and in parallel to vowel onsets (Oganian et al., 2023; Oganian & Chang, 2019). However, it is not known how these representations are affected by the stress status of individual vowels.

Here, we aim to identify the neural timescales for the effects of acoustic salience, lexical stress, and metrical predictability on neural responses to vowels in continuous speech. We capitalize on analytic approaches that allow time-resolved modeling of the neural signal as a function of speech input (Crosse et al., 2024; Holdgraf et al., 2017), disentangling between the effects of multiple concurrent properties of the speech signal. We hypothesize that acoustic salience and lexical stress are recognized online and affect early sensory processing of vowels. Furthermore, we hypothesize that these effects will be modulated by metrical predictability. For instance, the effect of acoustic saliency (i.e., vowel intensity) may be reduced when stress patterns are fully predictable. Alternatively, it is possible that meter and lexical stress are integrated only at later stages of vowel processing, with no effect on early sensory responses.

## Methods

### Stimulus materials: Seuss stories

The speech stimuli consisted of four children’s stories in two different meter conditions varying in predictability (Fig. 1a): metrically predictable (mp+) and metrically less predictable (mp-). The **mp+ condition** consisted of two Dr. Seuss (Scholastic Books) stories: “And to Think That I Saw It on Mulberry Street” and “McElligot’s Pool”. For the **mp-condition**, two modified versions of these stories were recorded. The modified versions were designed to exhibit less metrical patterns, deviating from Dr. Seuss’s characteristic *anapestic tetrameter* (that is two unstressed syllables followed by a stressed syllable) and emulating informal narrative styles, approximating a speech-like rhythm. Semantic content, syntactic structures, and sentence durations were maintained to align with the original texts. The speaker was instructed to maintain the metrical pattern of the regular stories, but to speak as naturally as possible. To ensure comparable syllable rates across stories, an experimenter monitored the speakers’ speech rate with the help of a metronome (set to 72bpm), which was not audible to the speaker.

**Figure 1.**
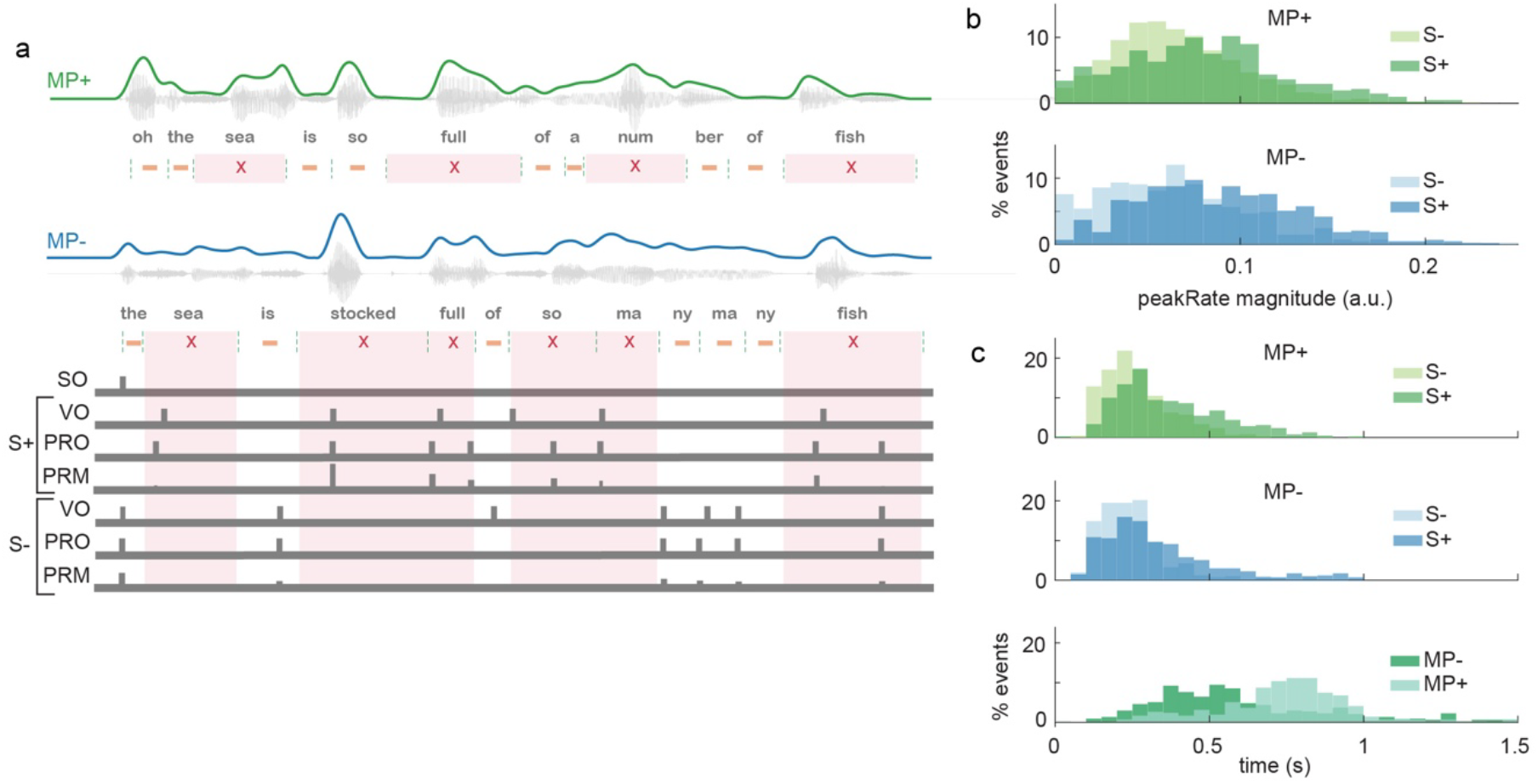
**a**. Example stimuli and design matrix. The acoustic waveform, amplitude envelope (green/blue), syllable boundaries (dashed lines) and stress pattern (x for stressed, - for unstressed) are shown for two example sentences in the MP+ (top) and MP- (bottom) conditions. In the MP+ condition, a stressed syllable is always followed by two unstressed syllables (anapestic tetrameter), for the MP-condition this meter is disrupted. For the MP-condition (bottom), the design matrix for the Temporal Response Function analysis is also shown. Sentence Onset is coded as a single predictor across MP conditions. Vowel onset (VO), peakRate onset (PRO) and peakRate magnitude (PRM) are coded separately for stressed and unstressed syllables, resulting in 13 predictors. **b**. Acoustic analysis of peakRate onset latencies and magnitudes in the stimulus materials. **c**. Top: Distribution of syllable durations (indexed as time intervals between peakRate onsets) by stress and meter condition. Bottom: Distribution of time intervals between stressed syllables (as indexed by peakRate onsets) by meter conditions.

### Stimulus annotation

#### Speech segmentation

The audio recordings were time-aligned to their respective orthographic transcriptions using the Montreal Forced Aligner (McAuliffe et al., 2017) (Version 3.9). Prior to alignment, all audio files were downsampled to 16 kHz to match the acoustic model’s feature generation specifications. Alignment employed the pretrained english_us_arpa acoustic model and its associated pronunciation dictionary. For any out-of-vocabulary word, phonetic transcriptions were generated using MFA’s Grapheme-to-Phoneme (G2P) model. The alignment process relied on a Hidden Markov Model-Gaussian Mixture Model (HMM-GMM) framework with speaker adaptation to improve boundary precision. Resulting phonemes and word-level timestamps were exported as Praat TextGrids for subsequent acoustic analysis (Boersma, 2001). To ensure temporal accuracy, all segment boundaries were manually inspected and refined by two independent coders (inter-rater reliability, cohen’s kappa > 0.8). The coders also inserted syllable-level boundaries where appropriate. All adjustments were based on detailed visual examination of the waveform and wide-band spectrogram, ensuring consistent and high-quality annotations.

#### Stress

Following boundary verification, the stress status of each syllable was manually annotated in Praat. A three-tier coding structure was employed to maintain high data quality and inter-rater consistency: A senior researcher established the annotation guidelines and performed gold-standard coding on a training set to calibrate the other raters. Two additional trained coders each processed 50% of the remaining dataset. To assess inter-rater reliability (IRR), a shared subset of 10% of the data was annotated by all three coders. Agreement was calculated using Cohen’s Kappa (or interrater reliability), with any discrepancies resolved through consensus discussions led by the master coder.

#### peakRate events

peakRates were computed following Oganian and Chang (2019), using publicly available code (https://github.com/yoganian/peakRate). In brief, we extracted the broad amplitude envelope of the speech signal, rectified it and low-pass filtered it at 10Hz, and calculated the derivative of this envelope. For each syllable, when more than one peakRate occurred, only the one closest to voicing onset was retained for analysis. We defined Inter-PeakRate intervals as the time from each peakRate to the subsequent peakRate, regardless of the stress status of the following syllable. We expected longer intervals in stressed syllables, reflecting longer stressed vowel durations.

#### Sentence onset

Sentence onset was manually annotated at the acoustic onset of each sentence.

### Data acquisition and preprocessing

Participants (n = 26, 21 females; age range 18-22 years, mean 18.8years, all English native speakers) passively listened to approx. 10 minutes of audio recordings consisting of two stories (one metric, one speech-like) while EEG data were collected. To ensure attentive listening, participants answered a set of multiple choice comprehension questions about the stories’ content after the experiment. Story pairings (MP- and MP+) and order of presentation were counterbalanced across participants to control for potential order effects. All participants provided informed written consent and participated in the study in exchange for study credit. The study protocol was approved by the Vanderbilt University IRB (#162002).

EEG data were continuously recorded using a 128-channel Hydrocel Geodesic Sensor Net (EGI, Eugene, OR, USA) with embedded Ag/AgCl electrodes. The net was placed on the scalp and connected to a Net Amps 200 amplifier via a Mac Pro computer. Electrode impedances were maintained below 50 kΩ, and data were referenced online to Cz. Vertical and horizontal electrooculograms (EOG) were also recorded to detect blinks and vertical eye movements. EEG and EOG signals were digitized at a sampling rate of 500 Hz.

EEG data were processed using EEGLAB (Delorme & Makeig, 2004) in Matlab R2024a (MathWorks, www.mathworks.com). Raw signals were initially downsampled to 250 Hz and high-pass filtered at 0.5 Hz. The PREP pipeline (Bigdely-Shamlo et al., 2015) was employed to identify and interpolate bad channels, apply robust average referencing, and remove 60 Hz line noise. Large transient artifacts were automatically detected and removed using Artifact Subspace Reconstruction (ASR, (Kothe & Makeig, 2013). Independent Component Analysis (ICA) was performed on a copy of the data downsampled to 100 Hz and high-pass filtered at 2 Hz to optimize algorithm efficiency. Resulting ICA weights were applied to the original data. ICLabel was used to automatically classify independent components, and components identified as eye movements with a probability exceeding 80% were systematically excluded.

### Data analysis

All analyses were performed in Matlab R2022b-R2025a (MathWorks, www.mathworks.com) using Fieldtrip (Oostenveld et al., 2011) and custom scripts.

#### Feature Temporal receptive field models

To analyze how neural responses to *peakRate* and *vowelOnset events* are modulated by meter and stress, we fitted temporal response functions (TRF) (Crosse et al., 2024; Holdgraf et al., 2017) using a 2 × 2 design with the factors *Meter* (metrically predictable mp+ vs. metrically less predictable mp-) and *Stress* (stressed vs. unstressed syllables). For each condition, *peakRates* were coded in two ways: as a *Binary* predictor marking the occurrence of a peakRate event, and as a *Magnitude* predictor indicating its strength. This allowed us to independently model the neural response to the occurrence of a peakRate event and to its magnitude, analogous to including separate categorical and parametric predictors in a multiple regression model. Finally, we included a *Sentence Onset* predictor to account for large responses typically observed at the beginning of speech following silence. In total, the predictor matrix comprised 13 predictors: eight for *peakRates* (binary and magnitude coding, separately for all combinations of stressed vs. unstressed syllables and mp+ vs. mp-meter conditions), four *vowelOnset* predictors (separately for all combinations of stressed vs. unstressed syllables and mp+ vs. mp-meter conditions) and one for sentence onset, irrespective of condition (Fig. 1a). Each subject’s data consisted of 82 trials corresponding to individual sentences, 41 in each meter condition. Mean sentence duration was 7 seconds (min 2 s, max 16 s). Model training was performed using five-fold leave-one-out cross-validation with an 80–20 train-test split (uniformly distributed across conditions). TRFs were estimated using time lags ranging from −100 to 600 ms with regularization parameters (λ) logarithmically spaced across 20 values from 1 to 10^7^. For each subject, λ was bootstrapped for each fold and channel separately, the optimal λ was selected based on maximal prediction accuracy. To ensure comparable beta weights across subjects and conditions, after fitting individual subject models, we computed generic TRF models, in which a single, shared regularization parameter (λ) was used. This “generic” λ was defined as the median λ values across all subjects. Model performance was then evaluated through *prediction accuracy*, as the Pearson’s (r) linear correlation coefficient between the actual and predicted data from the TRF model.

#### ROI Channel selection

We selected the 10 channels with the largest average TRF beta weight in a pre-defined time window of 0 to 200 ms post peakRate event. All reported analyses were performed in this region of interest (ROI).

#### Statistical analysis of TRF model beta weights

To assess the overall shape of neural responses to peakRate and vowelOnset events and the modulatory effect of peakRate magnitudes, we tested the average TRF weights against a pre-zero baseline, using a one-dimensional time-based cluster-based permutation tests (cluster-level alpha threshold of .05 in 1000 permutations) on TRFs averages across all channels in the ROI.

To assess statistical differences between conditions of interest in the TRF beta weight time courses, we conducted one-dimensional time-based cluster-based permutation tests (1000 iterations) for both main effects (Stress and Meter) and their interaction, separately for each peakRate (onset and magnitude) and vowel onset predictors. For this, effect TRFs were calculated for each effect separately and then tested against zero with the FieldTrip function ft_statistics_montecarlo (Oostenveld et al., 2011) and custom written matlab code. Effect TRFs were averaged across all channels in the ROI prior to testing, the cluster-level alpha threshold was set to .05. Effect TRFs were calculated as following:

Main effect (ME) of Stress = *MP+Stressed* + *MP-Stressed* - *MP+Unstressed* - *MP-Unstressed*

ME of Meter = *MP+Stressed* - *MP-Stressed* + *MP+Unstressed* - *MP-Unstressed*

Linear interaction of stress and meter (IA) = *MP+Stressed* - *MP-Stressed* - *MP+Unstressed* + *MP-Unstressed*

#### Latency analysis for TRF model beta weights

To evaluate differences in component latencies between conditions, we used a jack-knifing procedure (Kiesel et al., 2008), looking for the latency at which 80% of maximal deviation from 0 is reached within a pre-defined time-window. Time windows were defined based on the shape of the grand average TRF across conditions: Peak 1: 40 - 60 ms, Peak 2: 140 - 160 ms, Late negativity: 200 - 600 ms.

#### Inter-event phase coherence (IEPC)

To study neural phase locking to peakRate events, we computed inter-event phase coherence for each condition (Meter and Stress). A noncausal Butterworth bandpass filter was applied to extract 46 logarithmically spaced frequency bands ranging from 0.67 Hz to 15.2 Hz in .5-octave steps. The Hilbert transform was then used to obtain the instantaneous phase angle at each time point and frequency band for each condition. To reduce leakage of phase-locking between neighboring peakRate events, only events with no other event in the preceding 200 ms were included in the analysis. To assess statistical differences between conditions of interest in the IEPC analysis, we subtracted a pre-event baseline IEPC value for each condition and then tested for all main effects and interactions with two-dimensional time-frequency cluster-based permutation tests. These tests were performed using the FieldTrip function ft_statistics_montecarlo, averaging across all channels in the ROI, with a cluster-level alpha threshold of .05 in 1000 iterations.

## Results

### Acoustic analysis: peakRate is an acoustic cue to vowel onset and lexical stress

We first aimed to characterize the relationship between acoustic edges in the speech envelope (peakRate events) and syllabic stress. Based on prior literature (MacIntyre et al., 2022; Oganian & Chang, 2019), we hypothesized that peakRate events would mark both stressed and unstressed syllables, with larger peakRate magnitudes, a proxy of vowel intensity, and longer intervals to the following peakRate in stressed syllables. Overall, we found that each story contained between 790 and 1000 peakRate events, marking ∼90% of all syllables across stories. As expected, mean peakRate magnitudes were higher for stressed syllables (Fig. 1b: mp+: mean = 0.084 (SD = 0.044), mp-: 0.083 (SD = 0.045)) than for unstressed syllables (mp+: mean = 0.065 (SD = 0.039), mp-: 0.067 (SD = 0.039)) in both conditions (ME of stress: *F*(1, 3715) = 150.20, *p* < 0.001, ME of meter and interaction effect p > .5). In contrast, inter-peakRate intervals were not affected by stress or meter (all main effects and interactions p > .5, Figure 1c). This was likely because our child-directed stimulus materials contained fewer reductions than spontaneous speech (Fitzroy & Breen, 2020). Overall, this analysis confirmed peakRate as a reliable acoustic cue to syllable occurrence and lexical stress. The differences in peakRate magnitude between stressed and unstressed syllables also indicate that any analysis of stress effects must include peakRate magnitude as a covariate, to isolate categorical effects of stress perception from continuous effects of vowel intensity, reflected by peakRate magnitude.

### TRF model fit: Independent tracking of vowels and peakRate events

To isolate the independent evoked neural activity in response to vowels, peakRate events and sentence onsets as a function of stress and meter, we fitted temporal response function (TRF) models to the continuous speech data. Model prediction accuracies on a held-out test set were significantly above 0 (mean model r = 0.0385, t(25) = 6.65, p<0.001). Crucially, the full model (mean r = 0.39, SEM = 0.006) including all predictors outperformed a reduced model (mean r = 0.3, SEM = 0.004) that contained predictors for sentence onsets only (t(25) = 2.19, p = .04), confirming ongoing neural tracking of vowel onset and intensity.

### TRF time courses

TRF model weights revealed distinct timecourses of neural responses for sentence onsets, vowel onsets, peakRate events, and peakRate magnitudes (Fig. 2a-d). All four were localized to a central cluster, consistent with sources in bilateral auditory and speech cortices (Fig. 2a-d, right column). We first tested these TRF time courses against zero, to identify time windows with significant neural responses driven by each feature. Evoked responses to vowel onsets and peakRates showed an early bimodal shape, with peaks around 50 and 150 ms (Fig 2b-c). This early bimodal double-peak response was followed by a sustained negativity at 200 - 600 ms. This shape was distinctly different from the dynamics of responses to sentence onsets, which showed the N100-P200 complex (Figure 2a) post-event. A comparison of TRFs for peakRate event onsets and peakRate magnitudes (Fig. 2d) showed that the amplitude of all three key TRF responses (early bimodal and late sustained) scaled up with larger peakRate magnitudes, i.e., responses were stronger for acoustically more salient syllables. To further elucidate this effect, we reran the TRF model with peakRate values split in tertiles, each tertile modeled with separate binary peakRate predictors. The resulting TRFs (Fig. 2e) visualize the scaling of neural responses with peakRate magnitude.

**Figure 2.**
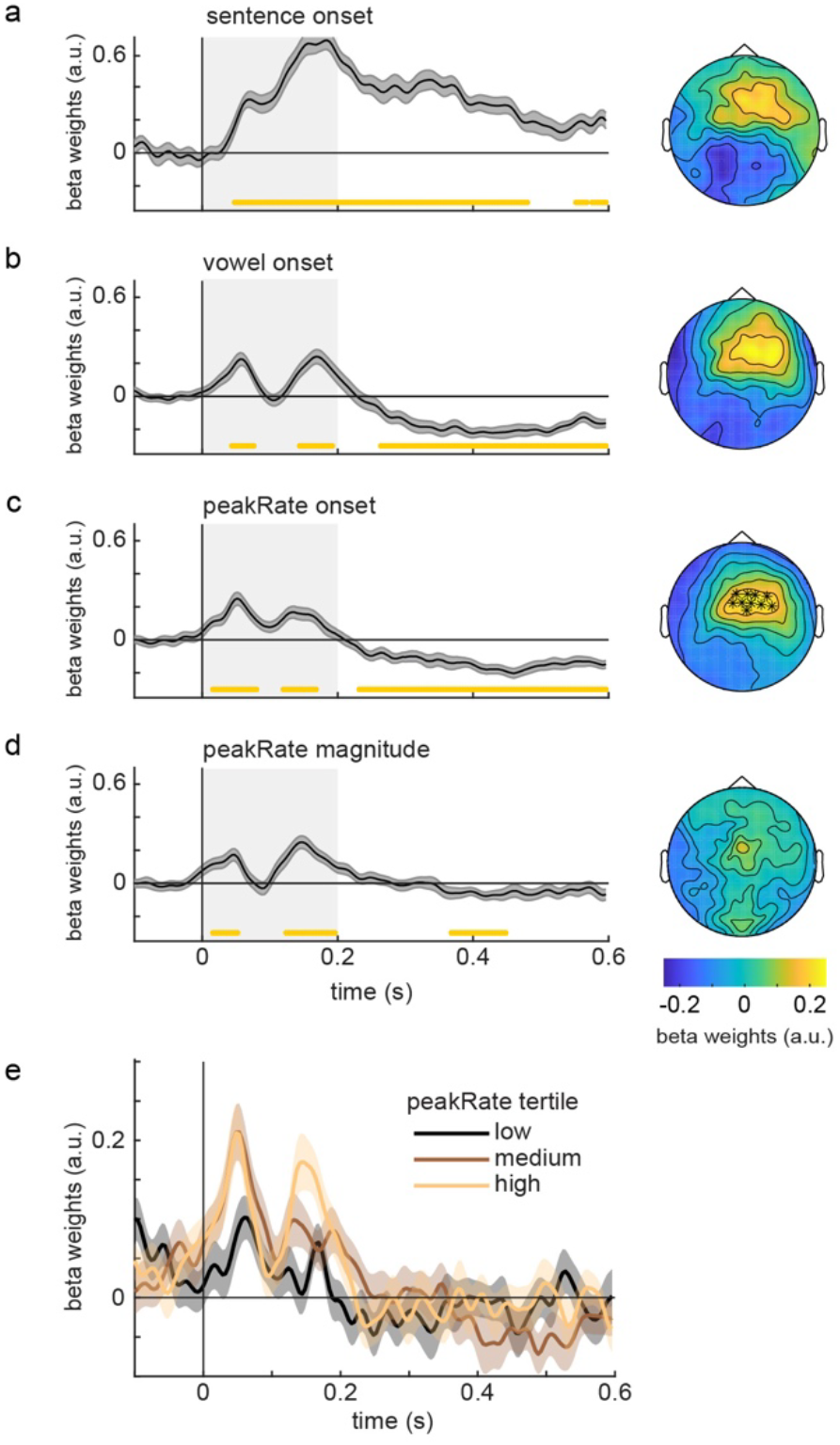
TRF weights by predictor. **a-d.** Timecourses and topographies for **a**. sentence onset TRFs; Grand average TRFs across conditions for each of the other model features: vowel onset (**b**.), binary peakRate events (**c**.) and peakRate magnitude (**d**.). Bottom yellow lines indicate time points significantly different from zero; boxes indicate the time window used for topographical maps (0 to 200 ms). **c**. stars in topography indicate channels included in ROI for all TRF analyses. **e**. peakRate TRFs by peakRate magnitude tertile show scaling of peakRate TRF by magnitude in the early timewindow (0 - 200ms).

Next, to understand how lexical stress and metrical speech context modulate neural tracking of vowels and peakRate events, we looked at the effects of stress and meter on TRF weights for each of the three predictor types (vowel onset, peakRate event, peakRate magnitude) separately.

### Vowel onsets: Responses integrate stress and meter information throughout processing

For vowel onsets, early sensory responses (first peak, ∼ 100 ms, Fig. 3a) were enhanced for unstressed as opposed to stressed syllables (cluster-based permutation p<0.05). In contrast, the late negativity to vowel onsets (starting at 400 ms) was stronger for stressed vowels than for unstressed vowels (ME Stress, cluster-based permutation p<0.05), and it was reduced when meter rendered syllable stress more predictable (ME Meter, cluster-based permutation p<0.05). Additionally, we observed a preparatory effect in the 50 ms prior to vowel onsets in the metrically less predictable condition (cluster-based permutation p<0.05). Taken together, this suggests that stress and its meter-based predictability affect vowel encoding throughout processing, with opposing effects at early and late stages. While early sensory processing is enhanced for unstressed vowels, late processing is stronger for stressed syllables and in metrically less predictable contexts. Furthermore, lack of metrical predictability appears to drive additional preparatory activity prior to vowel onset.

**Figure 3.**
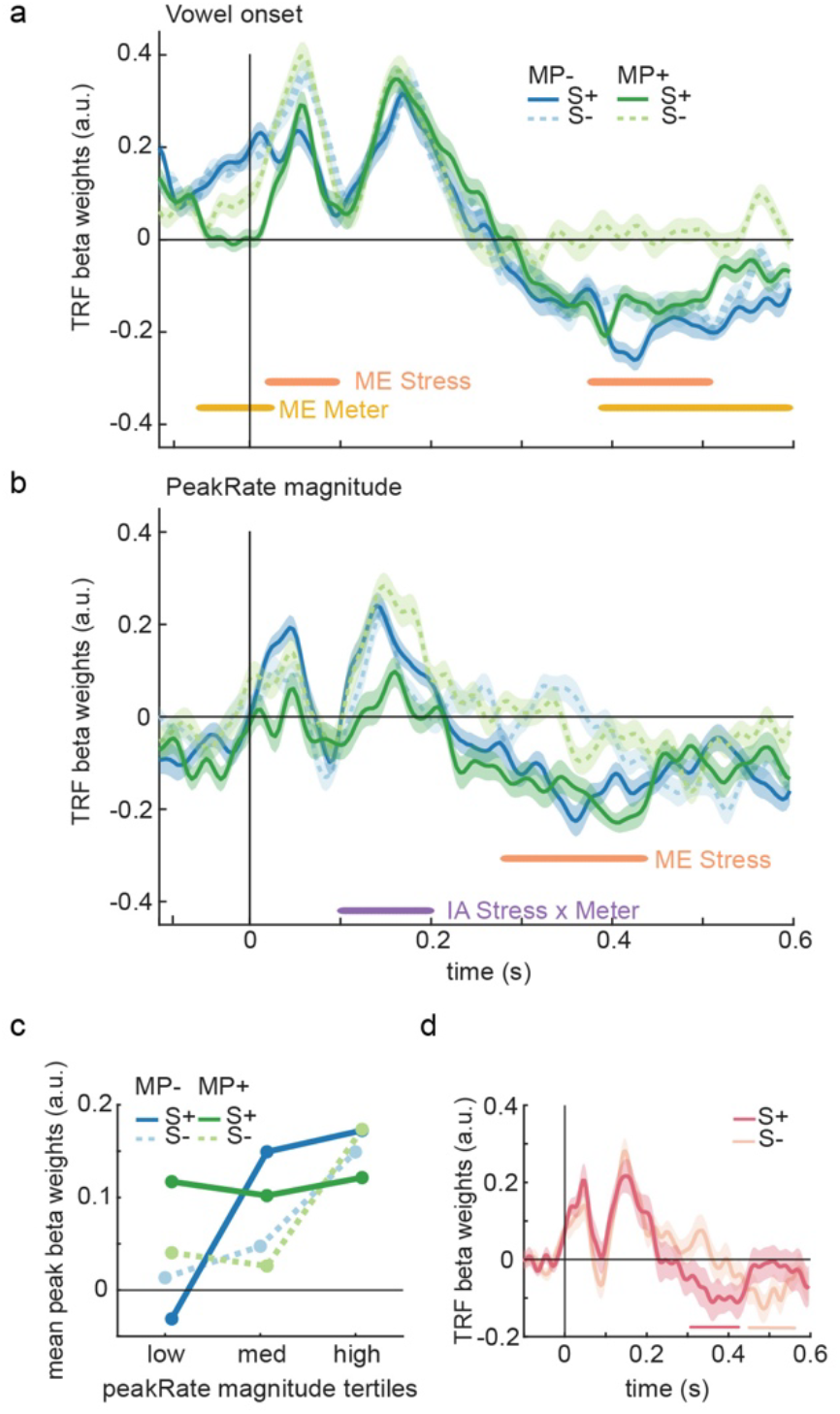
TRF weights by condition. **a**. Vowel onset. **b**. peakRate magnitude. **c**. Follow-up visualization of stress-meter interaction in the peakRate magnitude TRF for the 2nd early peak time window (125-175 ms). Neural responses to low and medium peakRate magnitudes are enhanced for stressed syllables in the MP+ condition. **d**. Follow-up visualization of the main effect of meter on the peakRate magnitude TRF. peakRate magnitude modulates magnitude of late negativity earlier for stressed (S+) than for unstressed (S-) syllables.

### PeakRate: Stress and meter reduce effects of event magnitude on neural tracking

For peakRate events, neither stress nor metrical predictability had a main effect on the overall pattern of neural responses. However, Early (prior to 100ms) responses to peakRate scaled with its magnitude (i.e., acoustic salience, cluster-based permutation p<0.05) irrespective of meter and stress (Fig. 3b). Starting from the second peak on, however, stress and metrical predictability interacted with the effect of peakRate magnitude. Specifically, in metrically less predictable stories, the magnitude of the second early peak (∼150ms, cluster-based permutation p<0.05) scaled with peakRate magnitude irrespective of stress. In contrast, in the metrically predictable story, peakRate magnitude modulated this peak in unstressed syllables but not in stressed syllables (cluster-based permutation p<0.05). To further qualify this effect, we again sorted peakRate values for each condition into tertiles (low, medium, high) and refit the model with binary predictors for each predictor. To understand the interaction of stress and meter effects in the second early peak, we extracted average TRF magnitudes for each condition and tertile in a 50-ms time window around this peak (Fig. 3c).

For peakRates with magnitudes in the medium tertiles, neural responses were larger for stressed than unstressed syllables, regardless of metrical predictability. However, the pattern differs for low and high tertiles. For low tertiles (i.e., low acoustic saliency), neural responses were large only when syllables were both stressed and metrically predictable; in all other cases, responses were near zero. At the other extreme, for high tertile peakRates, neural responses were consistently large, irrespective of metrical predictability or stress status. Thus, for stressed syllables in a metrically predictable context, acoustic saliency does not affect neural responses, leading to uniformly high response amplitudes. In contrast, when stress is not metrically predictable or when syllables are unstressed, neural response magnitudes in this early time window are determined by acoustic salience.

Finally, the scaling of the late negativity (∼ 400 ms) in TRF responses to peakRates with their magnitude were more pronounced for stressed syllables (Fig. 3b). A close inspection of this effect suggested that this effect may be due to latency differences. First, comparisons with the pre-zero baseline confirmed that TRFs for stressed and unstressed syllables both have the late negativity (Fig 3d). It occurred around 380ms for stressed syllables, but only at around 500ms for unstressed syllables. We thus used a jack-knifing procedure to estimate the latency of the late negativity at 80% of its maximum. This analysis confirmed a significantly earlier effect for stressed than for unstressed syllables (t = 8.63, p < 0.001).

In sum, the TRF analysis suggests three successive stages in the neural processing of lexical stress and meter. A preparatory stage, emerging around vowel onset, reflects preparatory activity in less predictable metrical contexts, suggesting anticipatory adjustments resulting from the lack of metrical expectancy. This is followed by an early sensory stage within the first 200 ms, encompassing the initial two response peaks. This stage shows heightened responses to acoustically less salient, unstressed vowels, and attenuated sensitivity to the acoustic saliency of expectedly stressed vowels. Finally, during the late sustained stage, processing is stronger for stressed syllables, especially when stress patterns are less predictable. Here, modulatory effects of acoustic saliency occur earlier for stressed than for unstressed syllables. Taken together, this shows that stress patterns and acoustic salience jointly modulate the temporal dynamics of feature integration throughout syllable processing.

### Inter Event Phase Coherence: Temporal consistency of response patterns depends on both stress and meter

Finally, we aimed to test how lexical stress and meter might affect neural phase locking to peakRate events, an acoustically defined landmark that is known to induce neural phase locking (Oganian et al., 2023). While TRFs and evoked response activity reflect the amplitude of neural responses, phase locking is a metric of precision in the timing of neural responses to these events. Particularly for peakRate, past work has shown that shape and occurrence timing of peakRate events in continuous speech determine spectral patterns in inter-event phase coherence (Oganian et al., 2023). Here we test how this pattern may be modulated by stress and meter. To this end, we calculated inter-event phase coherence (IEPC) between 0.67 and 15 Hz, aligned to the occurrence of peakRate events (Fig. 4). Across conditions, IEPC increased following peakRate events with spectral clusters roughly corresponding to the shape of the evoked response (around 6-10Hz), the average syllable rate (around 3.5-4 Hz) and the stress rate (around 1.3 Hz) in our stimulus materials, in line with prior findings (Oganian et al., 2023). Crucially, this pattern differed systematically between conditions (Fig. 4b-c). First, stress and meter affected IEPC at the syllable rate independently of each other, such that IEPC was lower for stressed syllables and in metrically predictable stories. That is, early sensory responses to peakRate were temporally more precise for unstressed than for stressed syllables. Second, IEPC in the 6-10 Hz range (evoked response) and in the 1.3 Hz range (stress rate) showed interaction effects of stress and meter. At 6-10 Hz, IEPC was stronger to stressed than to unstressed syllables when meter was less predictable, but this pattern was reversed when the meter became predictable. At the stress rate around 1.3 Hz, IEPC was higher for unstressed syllables in metrically less predictable stories, whereas in metrically predictable stories stress had no effect.

**Figure 4.**
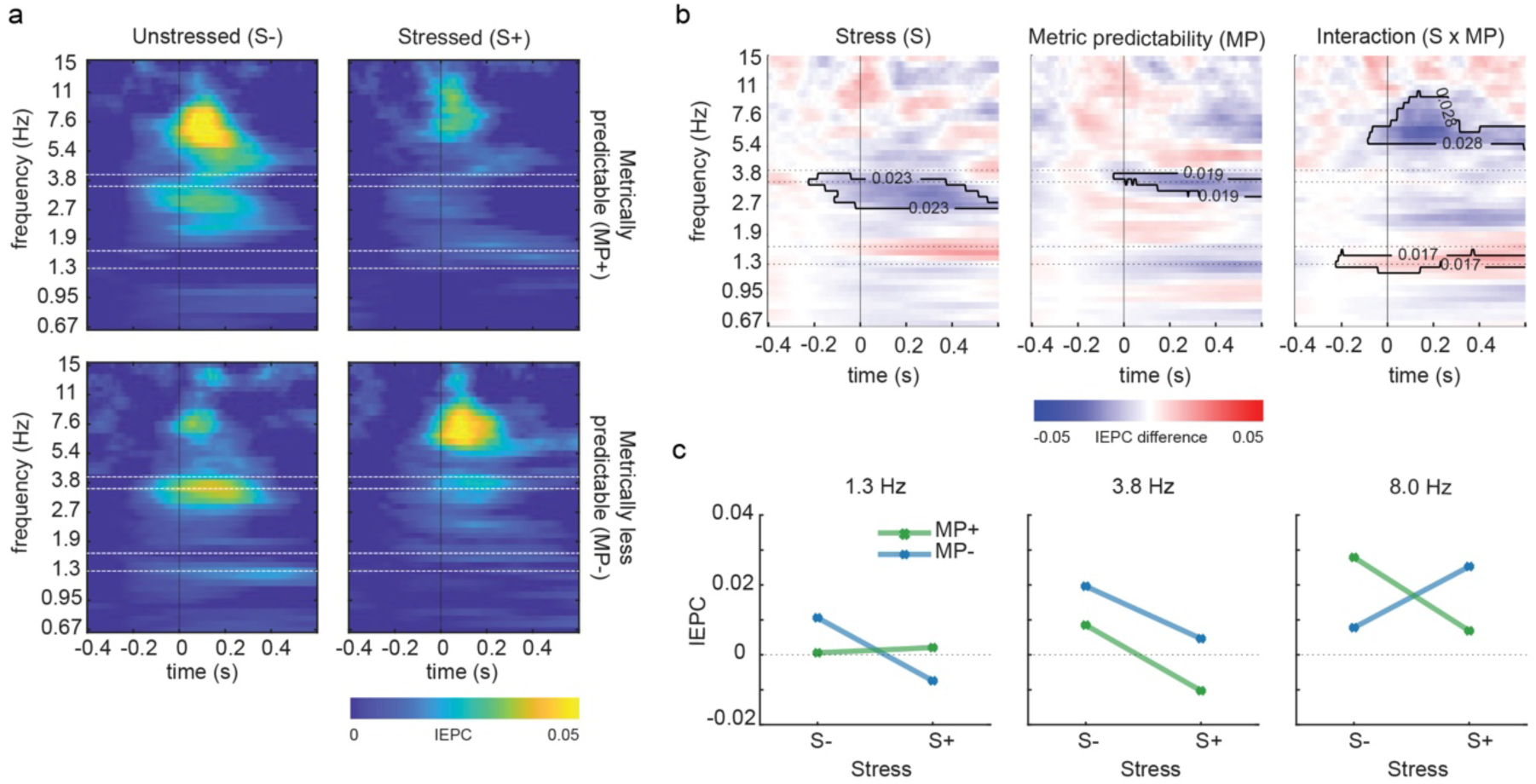
Inter-event phase coherence (IEPC) to peakRate events. **a**. Average IEPC to peakRate events by condition. Horizontal white lines delineate syllable frequency (around 3.8 Hz) and stressed syllable frequency (around 1.4 Hz) in the MP+ (top row) and MP-(bottom row) conditions, respectively. Neural phase locking is increased in three frequency bands, corresponding to the spectral shape of the evoked response (∼8Hz), the syllable rate (∼3.8 Hz) and the stress rate (∼1.3 Hz, respectively). **b**. Condition differences in IEPC show main effects of Stress and Meter ∼3.8 Hz, and interaction effects around 8 Hz and around 1.4 Hz. **c**. Average IEPC by condition for each of the three frequency ranges with significant effects of stress and meter conditions.

## Discussion

We isolate the effects of acoustic salience (i.e., magnitude of acoustic edges), linguistic salience (i.e., lexical stress) and the regularity of stress patterns (i.e., meter) on neural responses to continuous, storybook-like speech. We hypothesized that stress may be detected early on in a syllable with direct effects on processing of vowels and acoustic edges. In line with our expectations, temporal receptive field (TRF) analyses show effects of both lexical stress and meter throughout neural responses to vowels, including preparatory activity prior to vowel onset, early sensory responses (0-150ms), and late markers of integrational processes (300-600ms). Furthermore, we find that neural responses to low magnitude acoustic edges are enhanced within stressed vowels, even more so within a regular meter. Time-frequency analyses show that evoked responses to acoustic edges in the envelope are temporally more precise within unstressed than within stressed syllables, unlike previously assumed (Gross et al., 2013). In sum, our results demonstrate rapid integration of acoustic salience and lexical stress and speak in favor of parallel processing of acoustic, segmental and metrical information during speech perception.

Linguistic and psycholinguistic literature have debated the relative timing and relation between identification of individual speech sounds and lexical stress information. Behavioral (Cutler & Foss, 1977) and eye tracking evidence (Reinisch et al., 2010) points towards early recognition of stress concurrently with segmental information. However, electrophysiological evidence for this is missing, not least due to methodological issues associated with modeling of neural responses driven by complex speech acoustics. Here, we find that lexical stress affects neural responses to vowels within 100 ms post vowel onset. Critically, we find stronger early auditory evoked activity for unstressed than for stressed vowels. While this finding seemingly contradicts the dominant view that stressed vowels should drive stronger neural responses, it cannot be explained by acoustic differences between stressed and unstressed vowels, as those are accounted for in the TRF model. Rather, we propose that it might reflect additional effort in sensory encoding - possibly because recognition of vowel identity is harder for unstressed vowels (Calhoun, 2010). Alternatively, a reduced response to stressed vowels may be due to their higher predictability from preceding speech context. For instance, it has been found that the N100 auditory evoked component is reduced in regular and predictable contexts and enhanced for unexpected sounds (Schwartze et al., 2013).

Early integration of vowel and stress information is further supported by preparatory pre-vowel activity in metrically less predictable stories (MP-). Multiplex facilitatory effects of predictive processing have been observed across sensory domains, including language processing (Ryskin & Nieuwland, 2023). In speech, the benefits of prediction and expectation rely both on preparatory activity ahead of a stimulus (Goldstein et al., 2025), as well as ensuring ease of sensory processing (Ahissar et al., 2001) and later semantic integration (Frank et al., 2015). Here, we find that suprasegmental (stress) uncertainty can drive preparatory activity for individual speech segments, when their onset time - but not their stress status - can be expected based on preceding speech sounds. This shows that anticipatory processes are not confined to segmental and lexico-semantic processing but are pervasive across levels of speech analysis and provides evidence for rapid online integration between phonemic and prosodic structure during comprehension.

In addition to integration across linguistic levels, our findings support early integration of acoustic and linguistic and information. Scaling of neural responses to acoustic edges (peakRate events) with their magnitude was equivalent across stress and meter conditions in the first 100 ms. However, this scaling disappeared for stressed vowels in metrically predictable contexts. Rather, responses were strong even for edges of small magnitude. Neural responses to peakRate events are the primary source of speech envelope tracking (Oganian et al., 2023). The extent to which envelope tracking is a purely bottom-up auditory process or is subject to top-down modulation, such as intelligibility, has been subject to extensive research (Peelle et al., 2013). Our findings show that envelope tracking is supported by suprasegmental linguistic representations, enhancing tracking of smaller modulations in the envelope when they are expected to occur.

Early sensory responses to both vowels and peakRate events were followed by a late negativity around 300-500 ms after event onset, with topography and latency resembling the N400 component for semantically less expected words (Rabovsky et al., 2018). For vowels, this late negativity was stronger in metrically less predictable and stressed syllables. Numerically, however, both effects seem to reflect that this negativity was absent for unstressed vowels in metrically predictable contexts. Arguably, predictable unstressed vowels are easiest to identify as less informative and may thus be assigned less processing effort, similar to findings of an enhanced N400-like component to metrically unexpected words (Magne et al., 2007, 2016). While this past work was on violations of metric expectations, our stimuli did not contain a violation. Rather, they showed varying levels of metrical predictability. This speaks for continuous and graded predictions of upcoming stress patterns, akin to semantic expectations (Lau et al., 2008). Interestingly, for peakRates, this late effect was modulated by peakRate magnitude for both unstressed and stressed vowels, but at different latencies. Namely, it occurred later in unstressed than in stressed vowels, suggesting delayed integrative activity for unstressed but acoustically salient syllables. This speaks for a continuing role of acoustic salience even at later stages of syllable processing.

Complementary to our findings in the time domain, analyses of neural phase locking speak for increased temporal precision in neural tracking of peakRate events within unstressed vowels. It is known that evoked responses to peakRate events in the speech envelope directly map onto neural phase locking, with stronger phase locking to acoustically more salient events (Oganian et al., 2023). While TRF weights reflect the average response amplitude and temporal dynamics, neural phase-locking is a measure of temporal coherence in neural response shape and latency across trials. Here, we tested how stress and meter affect the temporal coherence of neural phase-locking to peakRate events. In our data, the phase-locked frequencies closely correspond to temporal modulation frequencies present in speech signal acoustics, lending additional support to an evoked response account of neural phase-locking to speech. Specifically, spectral peaks in phase-locking reflect the shape of the evoked response (6-8Hz), the syllable rate (∼4 Hz) and the stress rate (∼2 Hz) in our speech stimulus. In the metrically less predictable stories, phase-locking at 6-8 Hz, and thus temporal precision of evoked responses, was higher for stressed than for unstressed syllables. However, surprisingly, when the meter was predictable this pattern reversed, with higher temporal precision of responses for unstressed syllables. We propose that this reflects a shift in resource allocation: A predictable meter eases the processing of stressed syllables, allowing for more precise processing of unstressed syllables. In contrast, without metrical regularity the focus remains on the more informative stressed syllables. At the syllable rate, phase locking was evident in all conditions, except for stressed syllables in metrically predictable stories (where it, however, did appear at the stress rate). While further studies are necessary to assess the factors that determine this complex relationship, it does show that neural phase-locking depends not only on acoustic salience, but also on the interaction between linguistic salience and the predictability of the speech signal.

Ample evidence has shown that metrical regularity has beneficial effects on speech processing including acoustic (Moon et al., 2020), vowel (Pearson et al., 2021), and semantic information (Rothermich et al., 2012). Behaviorally, beneficial effects of metrical regularity have been documented for infant-directed materials and for adults (Cox et al., 2022; Cutler, A., 1991; Lenc et al., 2023; Pérez-Navarro et al., 2022). The present findings extend this literature by providing insight into the neural mechanisms underlying this facilitation both in early sensory processing and in later integration. Future research could examine whether atypical neural processing of metrical predictability contributes to observed behavioral deficits in language processing, such as those associated with developmental language disorders (Lense et al., 2021) and aging, and further explore its potential role in supporting language acquisition across development.

## Conclusion

We provide evidence for concurrent and rapid neural analysis and integration of acoustic, segmental (vowel) and suprasegmental (stress) patterns in continuous speech. We find that the current speech input is interpreted in context of suprasegmental metrical regularities and offer a possible neural basis for the beneficial effects of metrical regularities in speech processing.

## Author contributions

CZ: conceptualization, formal analysis, data curation, writing and editing, visualization. EB: formal analysis, editing. KK: conceptualization, formal analysis, data curation, editing. SP, JL, KD, KK, BM: Methodology, formal analysis, software. ML: conceptualization. SN: conceptualization, methodology, data curation, editing. RLG: conceptualization, methodology, software, investigation, resources, data curation, editing, supervision, project administration, funding acquisition. CM: conceptualization, methodology, software, investigation, resources, data curation, formal analysis, writing and editing, supervision, project administration, funding acquisition. YO: conceptualization, methodology, data curation, formal analysis, visualization, writing and editing, supervision, project administration, funding acquisition.

## Acknowledgements

CZ was supported by the German Academic Exchange Service (DAAD, Research Stays for University Academics and Scientists, Grant no. 57693448) and IBRO. EB was supported by the German Academic Exchange Service (DAAD Research Grants - Short-Term Grant no. 91901012). YO was supported by the German Research Council (DFG OG 105/4-1) and research group funding from the Werner Reichardt Center for Integrative Neuroscience (CIN).

## Notes

### Competing Interest Statement

The authors have declared no competing interest.

